# Ionic strength modulates HU protein-induced DNA supercoiling

**DOI:** 10.1101/2021.10.14.464438

**Authors:** Alexander Zhang, Yan Yan, Fenfei Leng, David Dunlap, Laura Finzi

**Affiliations:** Dept. of Physics, Emory University, 400 Dowman Dr., Atlanta, GA, 30322; Department of Chemistry and Biochemistry, Biomolecular Sciences Institute, Florida International University, 11200 SW 8th St., Miami, FL 33199

## Abstract

The histone-like protein from *E. coli* strain U93 (HU) is an abundant nucleoid-associated protein that contributes to the compaction of the bacterial genome as well as to the regulation of many of its transactions. Despite many years of investigations, the way and extent to which HU binding alters the DNA double helix and/or generates hierarchical structures using DNA as a scaffold is not completely understood. Here we combined single-molecule magnetic measurements with circular dichroism studies to monitor structural changes in the DNA-HU fiber as HU concentration was increased from 0 to 1000 nM under low and physiological monovalent salt conditions. We confirmed that DNA compaction correlated with HU concentration in a biphasic manner but DNA unwinding varied monotonically with HU concentration in 100 mM KCl. Instead, in more physiological 200 mM salt conditions, DNA compaction was monotonic while HU-induced DNA unwinding was negligible. Differential compaction and unwinding of DNA may be part of the response of bacteria to large variations in salt concentrations.

## Introduction

The concentration of total protein in the bacterial cytoplasm can rise to several milligrams per milliliter in exponentially growing cultures and remain above 6 mg/ml in stationary phase cultures [1]. A group of twelve nucleoid associated proteins constitute a significant fraction of this total, and two, the histone-like protein from *E. coli* strain U93 (HU) and the factor of inversion stimulation (Fis), are particularly prominent reaching up to 60,000 monomers/cell, approximately 60 μM, during exponential growth depending on the medium. These concentrations decline several-fold in stationary phase in which other proteins, DNA-binding proteins from starved cells (Dps) and phage integration host factor (IHF), become major structural components. Most nucleoid proteins oligomerize, and in addition, the existence of different isoforms can generate homo-oligomeric and hetero-oligomeric species with distinct DNA-binding characteristics. Indeed, HU αα-homodimers are prevalent in the exponential growth phase, but αβ-heterodimers dominate the early and late stationary phases [2, 3].

Several of these dimers acutely bend and induce supercoiling upon binding to DNA. Crystallographic studies of IHF bound to DNA revealed sharp bending of the DNA where intercalated proline residues at the tips of two β-ribbon arms kink the DNA backbone [4]. These prolines are well conserved among bacterial homologues, and the *Anabaena* HU protein produced similar, if somewhat more flexible, kinks in co-crystals with nicked DNA [5]. Studies of HU binding to plasmids in the presence of topoisomerase to relax unconstrained superhelicity demonstrated that HU binding at a 1:1 protein:DNA ratio negatively supercoiled DNA by −8 turns per 290 bp [6]. However, recent crystal structures of *E. coli* HU produced distinct complexes with DNA binding through a “phosphate lock” formed between G46 and K83, with no significant kinking of the DNA and little superhelical change in the DNA [7]. In this binding mode the two β-ribbon arms may interact with neighboring DNA molecules to form networks and bring about efficient nucleoid compaction in rapidly growing bacteria.

These disparate results may arise from different ionic strengths used in the respective experiments. Broyles and Pettijohn [6] used 50 mM NaCl for negative supercoiling measurements, while Hammel *et al*. [7] crystallized HU-DNA complexes in 200 mM CaCl_2_ or NH_4_F. Hammel *et al*. confirmed their crystallographic results with SAXS data on HU-DNA complexes in similar salt conditions, 150 mM NaCl. Matching the conditions is critical. Single molecule studies show that ionic strength determines the binding affinity and extent of compaction induced by HU [8, 9]. At low (40 – 100 mM NaCl) salt concentrations, titrating HU first compacts DNA up to 50-100 nM, but the DNA extends beyond the original length at 300 nM HU. In higher (150 mM NaCl) salt, HU progressively compacted DNA with no re-extension at higher HU concentrations, but HU did not compact DNA at all in 300 mM NaCl. Titrating with the *B. stearothermophilus* HU homologue (BstHU) in 100 mM KCl also progressively compacted DNA up to intermediate concentrations, and additional BstHU re-extended the DNA [10]. Either 2 or 200 nM HU negatively supercoiled the DNA under low tension in 100 but not in 200 mM KCl.

Other co-crystals of *E. coli* HU–DNA revealed octameric arrangements of monomers, including a left-handed template with the proper spacing for contacting a DNA molecule wrapped as a negatively supercoiled solenoid [11]. The authors also reported that increasing concentrations of HU progressively increased the negative supercoiling of plasmids incubated in 20 mM NaCl.

These results suggest that ionic strength modulates the binding mode of HU proteins, affecting the interplay between DNA compaction and supercoiling, the structure of the HU-DNA fiber and its functions. To investigate these macromolecular interactions, single DNA molecules were incubated with E. coli HUαβ and supercoiled to varying degrees with magnetic tweezers to measure HU-induced compaction and supercoiling. At 100 mM KCl, as the *E. coli* HU concentration was titrated, DNA molecules first compacted and then re-extended. Notably, while compaction induced by HU reversed at higher concentrations, HU-induced unwinding increased monotonically. Similarly, to what previously seen with BstHU, high ionic strength, 200 mM KCl, diminished compaction and negative supercoiling. Ionic strength appears to determine the binding mode of HU giving the protein broad activity while the helical form of the DNA remains unchanged as determined by circular dichroism spectra. This study highlights how the compaction and unwinding functions of HU are decoupled at low ionic strength such that a certain level of compaction corresponds to two different degrees of unwinding. The interaction simplifies at higher ionic strength, in which HU more mildly compacts DNA with little unwinding.

## Materials and Methods

### DNA tethers

The 3352 bp central fragment was produced by PCR with Taq DNA polymerase (New England BioLabs, Ipswich, MA, USA) from the template plasmid pYY_I1_400_BstEII[12] with forward (5-atgttcGGGCCCagtaacccgtatcgtgagc) and reverse (5’-acaacaCCCGGGatcatgtaactcgccttg) primers (Eurofins Genomics, Louisville, KY) including ApaI or XmaI restriction sites to generate overhangs for ligation. One end was ligated to a ∼100 bp digoxigenin-labeled fragments created by XmaI restriction of a 196 bp PCR amplicon of plasmid pUC19 produced with dATP, dCTP, dGTP, dTTP (Fermentas-Thermo Fisher Scientific Inc., Pittsburgh, PA, USA) and digoxigenin-11-dUTP (Roche Life Science, Indianapolis, IN, USA) in a molar ratio of 1:1:1:0.7:0.3. The other end was ligated to a ∼190 bp biotin-labeled fragments created similarly by ApaI restriction of a 375 bp PCR amplicon of plasmid pBluKSP containing biotin-11-dUTP (Invitrogen, Life Technologies, Grand Island, NY, USA). All fragments were purified with PCR purification kits (Qiagen, Germantown, MD, USA), and ligated with T4 ligase (New England BioLabs, Ipswich, MA, USA).

### Microchamber assembly

A microchamber with a volume of ∼15 uL with an inlet and an outlet was constructed with two glass coverslips (Fisherbrand, Thermo Fisher Scientific, Waltham, MA, USA) separated by a parafilm gasket and heat-sealed together [13]. DNA with digoxigenin at one end was attached to the anti-digoxigenin (Roche Life Science, Indianapolis, IN, USA) coated-coverslips, and a 1 um-diameter, streptavidin-coated paramagnetic bead (Dynabead MyOne Streptavidin T1, Invitrogen, Life Technologies, Grand Island, NY, USA) was tethered to the opposite, biotin-labeled end. The buffer solution for magnetic tweezing experiments was: 10 mM Tris-HCl (pH 7.4), 100 or 200 mM KCl, 5% DMSO, 0.1 mM EDTA, 0.2 mg/mL α-casein, and 0.2 mM dithiothreitol.

### Magnetic tweezing

A magnetic tweezer (MT) was used to supercoil DNA and to assay the dynamics of HU-mediated conformational changes in the DNA tethers. Permanent magnets were translated along and rotated about the optical axis of the microscope to vary the strength and orientation of the magnetic field in the microchamber [14, 15]. Intact, single DNA tethers were identified through extension vs. twist measurements [16]. Under tensions of approximately 0.25 pN, unwinding or winding causes torque in the DNA tether to increase until the buckling transition point is reached. At this point the first plectoneme is formed. Further twisting produces additional plectonemes which smoothly reduce the DNA extension. In contrast, nicked DNA swivels about single bonds to relax any applied torsion, fails to form plectonemes, and the extension does not decrease.

Tension for all experiments with HU protein was maintained at ∼0.25 pN as the HU protein was serially diluted with buffer. Extension versus twist measurements were recorded after ∼1 minute incubation with successive additions of increasingly concentrated HU protein up to 1000 nM, a concentration for which the compaction induced by HU protein reaches a plateau as reported in previous tethered particle motion experiments [17].

### Compaction and shift analysis

The compaction and shift of the curves recorded in the presence of different concentrations of HU protein with respect to that without protein was determined by fitting points above 50% extension on the respective curves with parabolas and comparing the vertices (Figure S1). The lengths of DNA tethers varied slightly due to different fractions of the tails being attached to the microchamber, or bead, surface [12] and the maximum supercoiling and compaction induced by the HU protein was proportional to the length of the DNA. Therefore, to plot an overlay of the entire data set, the shift and compaction determined for each DNA tether was scaled according to the length of the tether with no added twist in the absence of HU protein.

### CD Spectra

CD spectra were recorded at room temperature using a Jasco 1500 with a 1 cm path length cuvette filled with 150 ul of 7.3 or 10 μM DNA and varying concentrations of HU in 10 mM Tris-HCl pH 7.4, 100 or 200 mM KCl, and 0.1 mM EDTA. Background spectra with buffer only were subtracted from spectra for HU+DNA samples.

## Results

Magnetic tweezers are a single molecule technique which allows manipulation of the tensional and torsional state of biomacromolecules [18-21]. A measurement of the DNA extension as a function of negative or positive turns imparted to the molecule produces curves as shown in Figure 1. For tensions below about 0.4 pN, extension vs. twist curves are symmetrical and peak when the DNA is torsionally relaxed.

**Figure 1.**
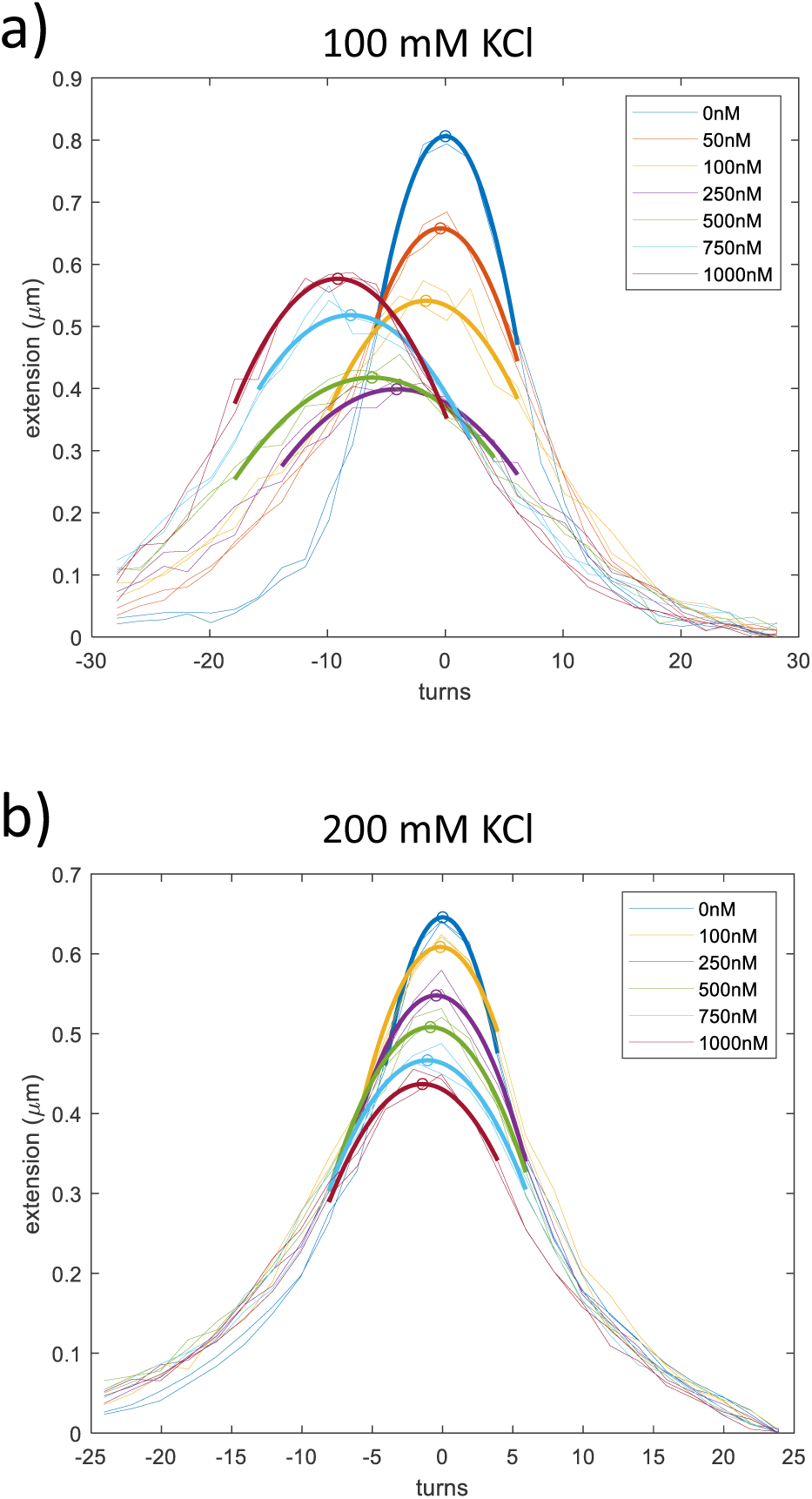
HU protein compacts and negatively supercoils DNA. (A) Titrating the concentration of the HU protein in 100 mM KCl from 0 to 250 nM progressively negatively supercoiled DNA tethers and compacted them. Additional HU up to a saturating amount further negatively supercoiled the DNA but reversed about half of the previous compaction. (B) In contrast, titrating the concentration of HU in 200 mM KCl progressively negatively supercoiled and compacted the DNA tether to much lesser degrees, and high HU concentrations merely augmented and did not partially reverse the compaction previously induced at lower concentrations.

Extension vs. twist curves were recorded for DNA under 0.25 pN tension, while titrating the HU protein from 0 to 1 μM at 100 or 200 mM KCl salt concentrations. Figure 1a shows a representative HU titration of a 3342 bp DNA tether in 100 mM KCl (for a gallery of titrations see Figure S2). In 100 mM KCl, the maximum extension of the DNA tethers initially dropped as HU was increased from 0 to 250 nM but subsequently increased as HU concentration was raised to 500, 750 and finally 1000 nM. As reported previously, increasing the HU concentration initially compacts DNA but subsequently reduces the compaction. During the titration, the peak of the hat curve also shifted indicating a progressive DNA unwinding as the HU concentration increased. Remarkably, HU-induced compaction of the DNA tether in 100 mM KCl was biphasic while unwinding was monotonic. Changing the KCl concentration altered this behavior. Figure 1b shows the change in DNA extension versus twist curves during a similar HU titration in 200 mM KCl (for a gallery of titrations see Figure S3). Compared to 100 mM KCl, HU in 200 mM KCl compacts DNA less and negligibly unwinds DNA. Figures 2a & 2b, where the maximum DNA tether extension is plotted as a function of HU-induced supercoiling, summarize these observations. Notably, at the more physiologically relevant concentration of 200 mM KCl [22, 23], HU-induced supercoiling rarely exceeds −2 turns (Sigma = 0.6%).

**Figure 2.**
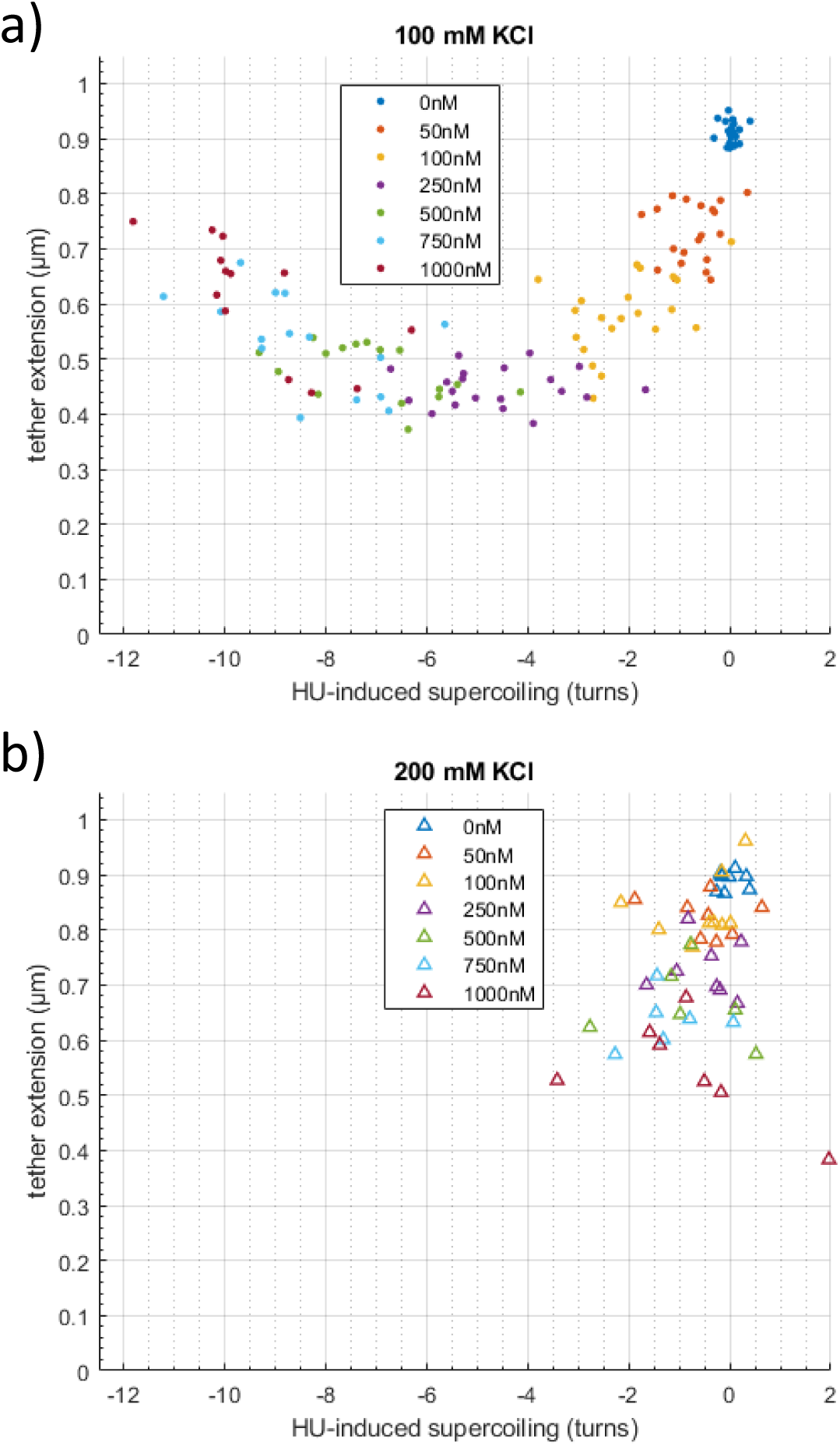
The HU protein compacts and supercoils DNA. A) Titrating the concentration of the HU protein in 100 mM KCl from 0 to 250 nM progressively negatively twisted DNA tethers and compacted them Higher concentrations of HU up to a saturating amount continued to negatively supercoil the DNA and reversed about half of the previous compaction. B) In contrast, titrating the concentration of HU in 200 mM KCl progressively negatively supercoiled and compacted the DNA tether to a much lesser degrees, and high HU concentrations merely augmented the compaction previously induced.

The stark difference between the HU effect on DNA compaction and unwinding at the two different ionic strength conditions, raised the hypothesis that, besides weakening electrostatic interactions between HU and DNA, higher monovalent cation concentrations may change the mode of binding of HU to DNA. Alternatively, HU-induced supercoiling may differentially partition as writhe at 100 mM KCl and twist at 200 mM KCl. Circular dichroism spectra are useful for revealing such changes in the helical structure of DNA [24]. Indeed, Johnson et al., combining experiments with simulations, argued that a decrease in DNA ellipticity (CD amplitude) is connected to an increase in the winding angle of the double helix, or a decrease in the twist [25]. Thus, circular dichroism (CD) spectra were recorded using 3352 bp-long DNA and different concentrations of HU (50, 250 and 1000 nM). Figure 3 shows that HU did not significantly change the CD spectra in the range from 240 to 300 nm, which is sensitive to DNA structure, at either salt concentration.

**Figure 3.**
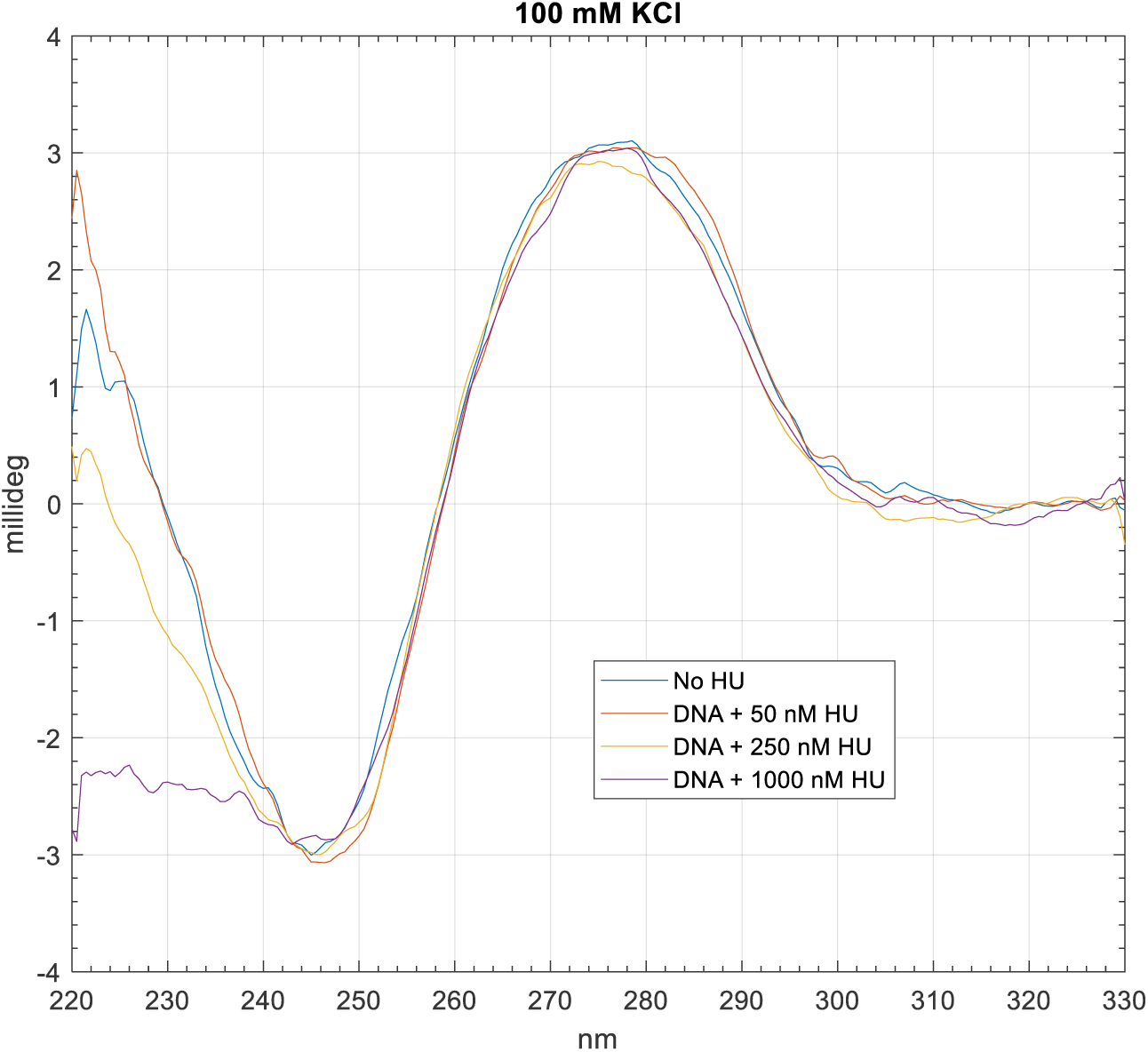
In CD spectra recorded for 7.3 uM of the DNA tether fragment in buffers containing 100 mM KCl and several concentrations of HU, there is no change in the range that reports on the DNA structure, 245-300 nm. As the HU concentration increased, the CD signal became progressively more negative in the range from 220 −245 nm, where the protein signal dominates.

## Discussion

HU is thought mediate long-range interaction between distant domains in the bacterial nucleoid [26]. The measurements reported here, under mild, 0.25 pN, tension do not display such long-range, *cis* interactions that would produce loops, although. previous studies [22, 23] indicate that 200 mM KCl is likely to mimic physiological, ionic strength in *E. coli* while 100 mM KCl may be low. Thus, at physiologically relevant ionic strength, HU-induced DNA compaction and unwinding is not maximal, and the homeostatic level of HU-induced unwinding of *E. coli* chromatin, might be no greater than 1% (see Results and Figure 2b). This would be consistent with the results of these single molecule, *in vitro* experiments.

However, other *E. coli* proteins also regulate the compaction and function of the nucleoid, and their aggregate effect may produce greater unwinding and compaction. Also, HU might produce negative supercoiling in other ways, for example, by stabilizing the transient opening of bubbles, or interacting with other unwinding factors [27-29]. In the closed, circular bacterial genome, any unwinding of the double helix introduced by HU binding would cause compensatory, free (unconstrained by protein), positive supercoiling which then needs to be relaxed. To avoid that outcome, the amount of DNA unwinding by an abundant, architectural protein like HU may be limited at physiological salt concentrations. It may not be beneficial for the cell to have to cope with excessive accumulation of free positive supercoiling in the genome.

In agreement with previous reports, ionic strength determined HU-driven DNA compaction without significantly modifying DNA structure. In addition, these magnetic tweezing experiments indicate that HU-induced unwinding of DNA is salt dependent and may be minimal in physiological salt conditions.

## Acknowledgements

This work was supported by the National Institutes of Health [R01GM084070 to L.F., and 1R15GM109254-01A1 to F.L.]. Katie Whitcomb performed preliminary CD experiments.

---

**Fig S1.**
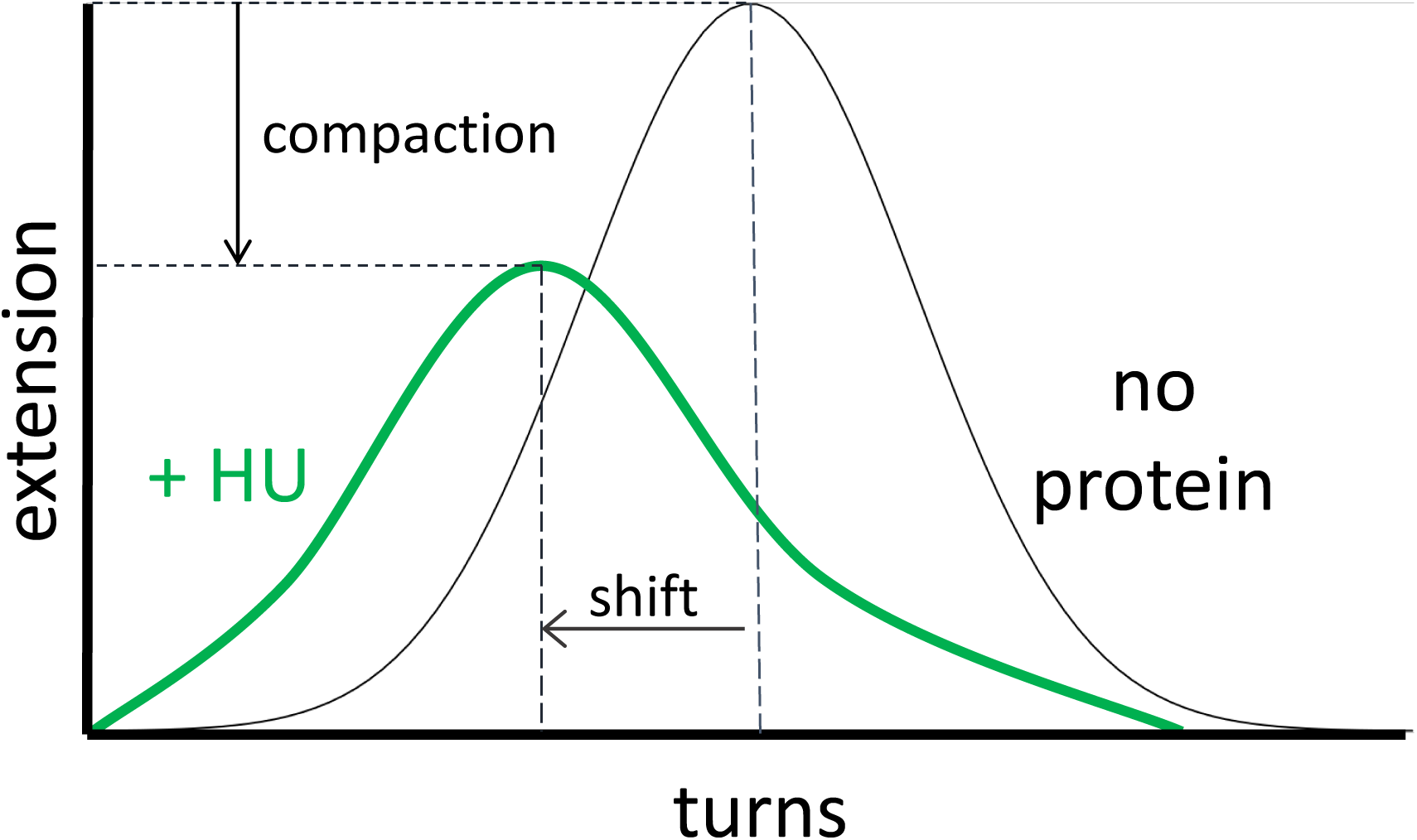
A schematic diagram of he analysis of extension versus twist curves produce by the titration of HU concentration. The vertex of each curve was compared to the vertex of the curve recorded for DNA without HU to determine how much compaction and supercoiling were induced by HU.

**Fig S2.**
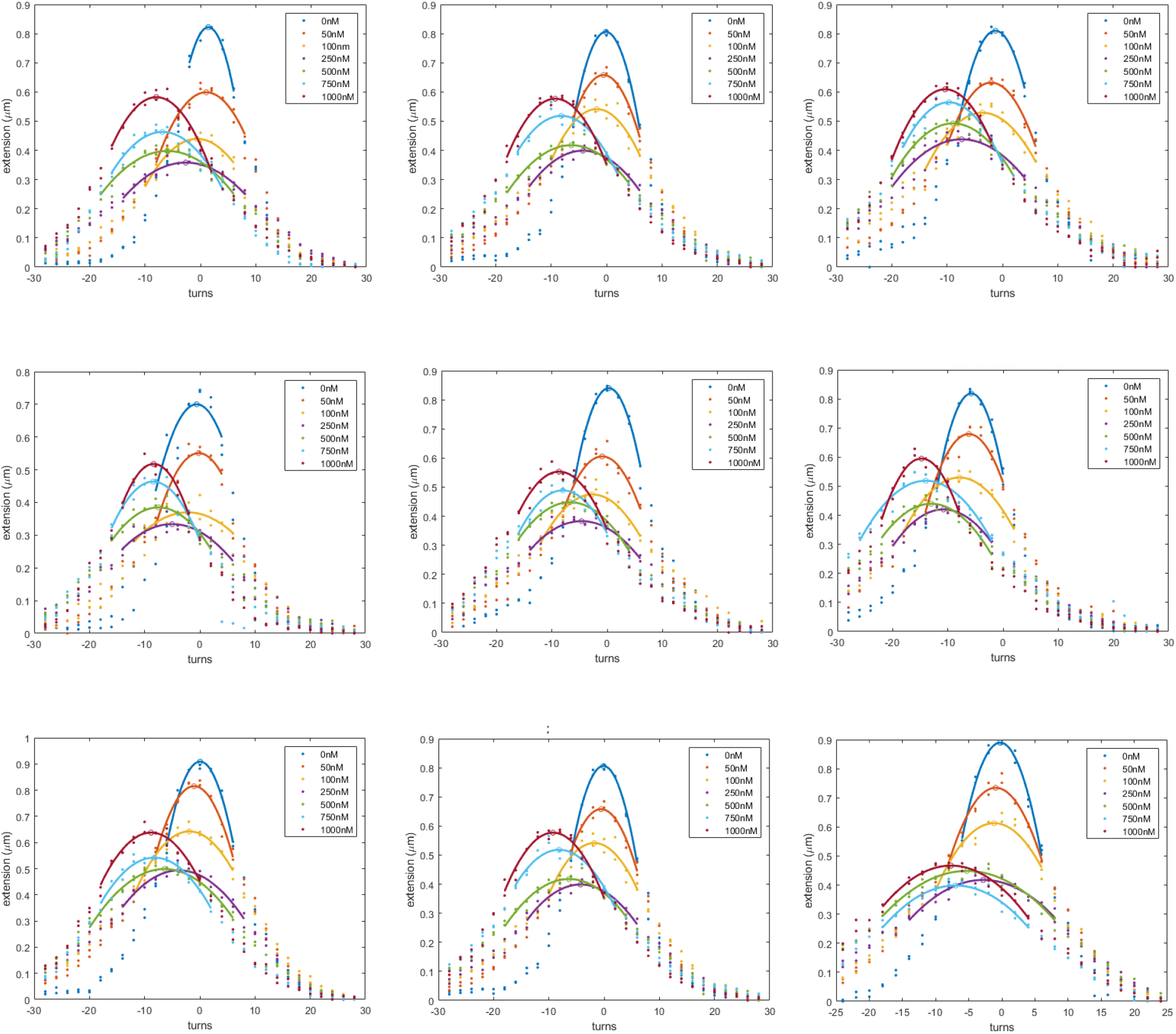
Significant compaction but negligible shift is obvious in examples of raw data from DNA tethers in 100 mM KCl with increasing concentrations of HU protein. Extension versus twist curves were recorded for 3352 bp DNA tethers stretched between digoxigenin-coated coverslips and streptavidin-coated MyOne beads with 0.25 pN of tension. Colors indicate HU concentrations. The data (dots) above half of the maximum height were fit with a parabola to establish the extension and HU protein-induced twist.

**Fig S3.**
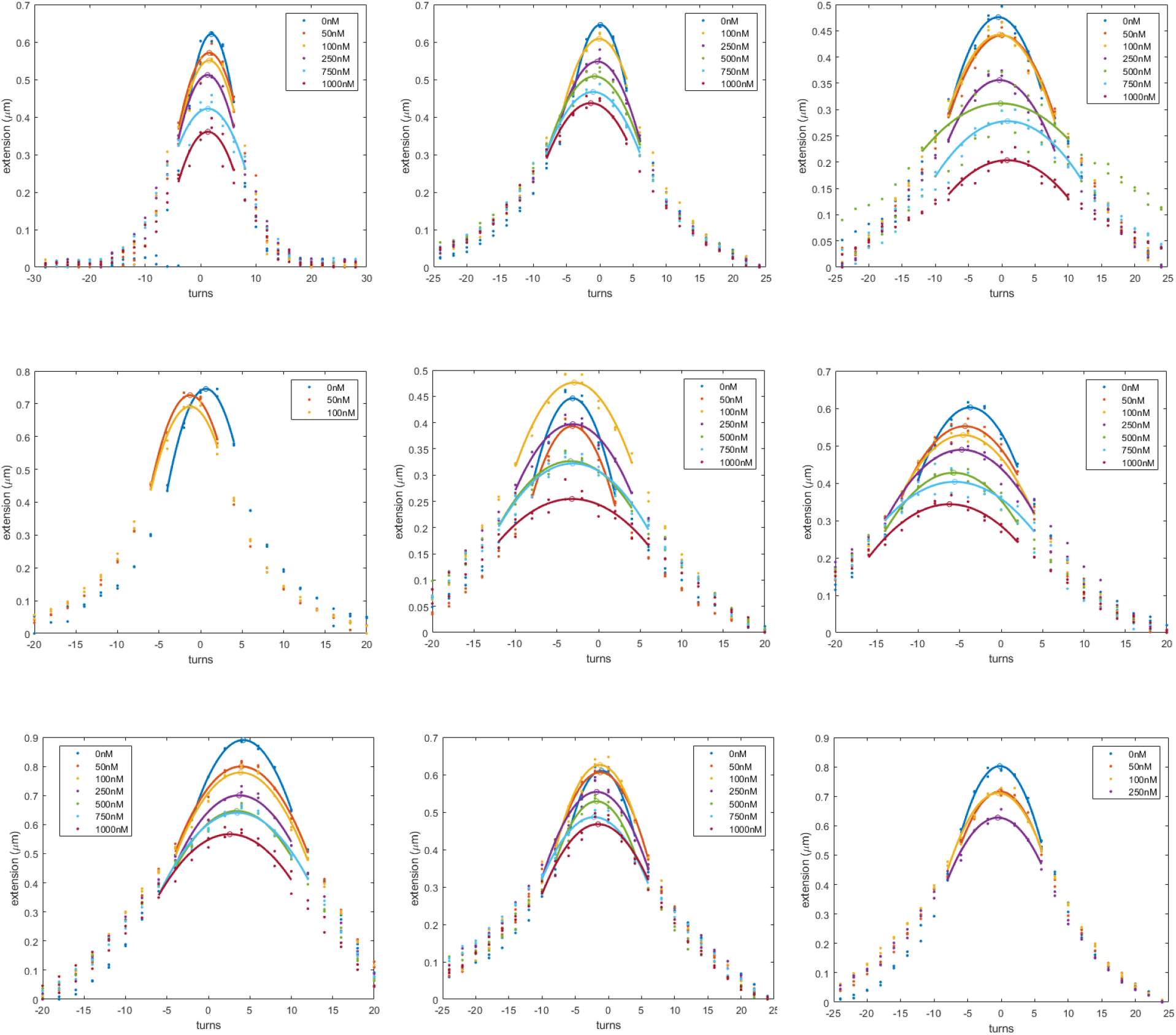
Significant compaction but negligible shift was obvious in raw data from DNA tethers in 200 mM KCl with increasing concentrations of HU protein. Extension versus twist curves were recorded for 3352 bp DNA tethers stretched between digoxigenin-coated coverslips and streptavidin-coated MyOne beads with 0.25 pN of tension. Colors indicate HU concentrations. The data (dots) above half of the maximum height were fit with a parabola to establish the extension and HU protein-induced twist.

## References

1. Ali Azam, T., et al., Growth Phase-Dependent Variation in Protein Composition of the Escherichia coli Nucleoid. Journal of Bacteriology, 1999. 181(20): p. 6361–6370.

2. Claret, L. and J. Rouviere-Yaniv, Variation in HU composition during growth of Escherichia coli: the heterodimer is required for long term survival. Journal of Molecular Biology, 1997. 273(1): p. 93–104.

3. Tanaka, H., et al., Properties of DNA-binding of HU heterotypic and homotypic dimers from Escherichia coli. J Biochem, 1993. 113(5): p. 568–72.

4. Rice, P.A., et al., Crystal structure of an IHF-DNA complex: a protein-induced DNA U-turn. Cell, 1996. 87(7): p. 1295–306.

5. Swinger, K.K., et al., Flexible DNA bending in HU-DNA cocrystal structures. Embo J, 2003. 22(14): p. 3749–3760.

6. Broyles, S.S. and D.E. Pettijohn, Interaction of the Escherichia coli HU protein with DNA: Evidence for formation of nucleosome-like structures with altered DNA helical pitch. Journal of Molecular Biology, 1986. 187(1): p. 47–60.

7. Hammel, M., et al., HU multimerization shift controls nucleoid compaction. Science Advances, 2016. 2(7): p. e1600650.

8. Xiao, B., R.C. Johnson, and J.F. Marko, Modulation of HU–DNA interactions by salt concentration and applied force. Nucleic Acids Research, 2010. 38(18): p. 6176–6185.

9. Lin, S.-N., R.T. Dame, and G.J.L. Wuite, Direct visualization of the effect of DNA structure and ionic conditions on HU–DNA interactions. Scientific Reports, 2021. 11(1): p. 18492.

10. Schnurr, B., C. Vorgias, and J. Stavans, Compaction and supercoiling of single, long DNA molecules by HU protein. Biophysical Reviews and Letters, 2006. 01(01): p. 29–44.

11. Guo, F. and S. Adhya, Spiral structure of Escherichia coli HUaβ provides foundation for DNA supercoiling. Proc Natl Acad Sci U S A, 2007. 104(11): p. 4309–14.

12. Yan, Y., et al., Protein-mediated looping of DNA under tension requires supercoiling. Nucleic Acids Research, 2018. 46(5): p. 2370–2379.

13. Kovari, D.T., et al., Tethered Particle Motion: An Easy Technique for Probing DNA Topology and Interactions with Transcription Factors. Methods in Molecular Biology, 2018. 1665: p. 317–340.

14. Strick, T.R., et al., The elasticity of a single supercoiled DNA molecule. Science, 1996. 271(5257): p. 1835–7.

15. Kovari, D.T., et al., Model-free 3D localization with precision estimates for brightfield-imaged particles. Optics Express, 2019. 27(21): p. 29875–29895.

16. Charvin, G., et al., Tracking topoisomerase activity at the single-molecule level. Annu Rev Biophys Biomol Struct, 2005. 34: p. 201–19.

17. Yan, Y., et al., Protein-mediated looping of DNA under tension requires supercoiling. Nucleic Acids Res, 2018. 46(5): p. 2370–2379.

18. Charvin, G., et al., Twisting DNA: single molecule studies. Contemporary Physics, 2004. 45(5): p. 383–403.

19. Finzi, L. and D.D. Dunlap, Single-molecule approaches to probe the structure, kinetics, and thermodynamics of nucleoprotein complexes that regulate transcription. J Biol Chem, 2010. 285(25): p. 18973–8.

20. Gosse, C. and V. Croquette, Magnetic tweezers: micromanipulation and force measurement at the molecular level. Biophys J, 2002. 82(6): p. 3314–29.

21. Piccolo, J.G., et al., Force spectroscopy with electromagnetic tweezers. Journal of Applied Physics, 2021. 130(13): p. 134702.

22. Schramke, H., et al., Revisiting regulation of potassium homeostasis in Escherichia coli: the connection to phosphate limitation. MicrobiologyOpen, 2017. 6(3): p. e00438.

23. Shabala, L., et al., Ion transport and osmotic adjustment in Escherichia coli in response to ionic and non-ionic osmotica. Environmental Microbiology, 2009. 11(1): p. 137–148.

24. Kypr, J., et al., Circular dichroism and conformational polymorphism of DNA. Nucleic Acids Res, 2009. 37(6): p. 1713–25.

25. Johnson, B.B., et al., Correlations between deoxyribonucleic acid structural parameters and calculated circular dichroism spectra. Biochemistry, 1981. 20(1): p. 73–8.

26. Lioy, V.S., et al., Multiscale Structuring of the E. coli Chromosome by Nucleoid-Associated and Condensin Proteins. Cell, 2018. 172(4): p. 771-783.e18.

27. Hwang, D.S. and A. Kornberg, Opening of the replication origin of Escherichia coli by DnaA protein with protein HU or IHF. J Biol Chem, 1992. 267(32): p. 23083–6.

28. Polaczek, P., K. Kwan, and J.L. Campbell, Unwinding of theEscherichia coliOrigin of Replication (oriC) Can Occur in the Absence of Initiation Proteins but Is Stabilized by DnaA and Histone-like Proteins IHF or HU. Plasmid, 1998. 39(1): p. 77–83.

29. Ryan, V.T., et al., IHF and HU stimulate assembly of pre-replication complexes at Escherichia coli oriC by two different mechanisms. Molecular Microbiology, 2002. 46(1): p. 113–124.

